# Simulations reveal that beta burst detection may inappropriately characterize the beta band

**DOI:** 10.1101/2023.12.15.571838

**Authors:** Zachary D Langford, Charles R E Wilson

## Abstract

In neurophysiological research, the traditional view of beta band activity as sustained oscillations is being reinterpreted as transient bursts. Bursts are characterized by a distinct wavelet shape, high amplitude, and, most importantly, brief temporal occurrence. The primary method for their detection relies on a threshold-based analysis of spectral power, and this presents two fundamental issues. First, the threshold selection is effectively arbitrary, being influenced by both temporally proximal and distal factors in the signal. Second, the method necessarily detects temporal events, as such it is susceptible to misidentifying sustained signals as transient bursts. To address these issues, this study systematically explores burst detection through simulations, shedding light on the method’s robustness across various scenarios. Although the method is effective in detecting transients in numerous cases, it can be overly sensitive, leading to spurious detections. Moreover, when applied to simulations featuring exclusively sustained events, the method frequently yields events exhibiting characteristics consistent with a transient burst interpretation. By simulating an average difference in power between experimental conditions, we illustrate how apparent burst rate differences between conditions can emerge even in the absence of actual burst rate disparities, and even in the absence of bursts. This capacity to produce misleading outcomes challenges the reinterpretation of sustained beta oscillations as transient bursts and prompts a critical reassessment of the existing literature.

**New and Noteworthy:** Neurophysiological research is experiencing a transformative shift in understanding beta band activity, moving away from the notion of sustained oscillations towards recognizing the significance of transient bursts. Here we show how the methods to detect such bursts are prone to spurious detections and can blur the distinction between sustained signals and transient bursts. Further, in realistic scenarios these methods can produce apparent behavioral associations where no such association exists.

## INTRODUCTION

Beta bursts are transient spectral events, occurring between 15 and 29 Hz, in neurophysiological recordings. Bursts are usually characterized as being of relatively high amplitude, occurring briefly (<300 ms), and having the distinct appearance of a wavelet shape (Sherman et al., 2016; Szul et al. 2022; Zich et al. 2023). Beta bursts have been linked to sensorimotor cortex during motor actions (Wessel 2020; Bonaiuto et al., 2021; Zich et al. 2023; Rayson et al., 2023) and sensory attention (Sherman et al., 2016), prefrontal cortex during working memory tasks (Lundqvist et al. 2018; Liljefors et al., 2023), and frontal cortex in cognitive control tasks (Langford, Procyk, & Wilson, 2023). However, despite their functional relevance, the precise nature and underlying mechanisms of beta bursts remain incompletely understood, and this may largely be determined by the methodologies employed to characterize them.

The most common method used to detect bursts is based on identifying a threshold (PT-method; also known as the p-episode method (Tal et al., 2020)), or cutoff point, in power - at which any value occurring above this threshold is considered a putative burst (e.g., Shin et al., 2017) if it fulfills a number of other criteria. The threshold is often chosen as the point (Shin et al., 2017) that maximizes the correlation between the single-trial area above the threshold and averaged single-trial power. This threshold is contended to be appropriate because it most fully accounts for the variance in single trial power (Shin et al., 2017), yet in of itself this contention makes the strong assumption that the transient bursts are the only factor contributing to single trial power. It has further become commonplace to employ a predetermined threshold, commonly set at six factors-of-medians (FOM), across a gamut of experimental designs, recording modalities, cortical locations, and species (e.g., Wessel et al. 2020; Morris et al., 2023; Errington, Woodman, & Schall, 2020; Law et al., 2022; and additionally in the gamma band: McKeon et al., 2023).

In this study, our aim was to conduct a comprehensive investigation into the performance of the PT-method using simulated data. This strategy is particularly useful because it enables the modeling of numerous plausible burst scenarios, allowing us insight into the ground truth of each, which is not attainable through neural data alone. We narrow our focus on addressing what we view as two unresolved issues. First, the establishment of an optimal cutoff using correlation maximization appears to harbor an element of arbitrariness. This stems from the fact that the identification of individual bursts hinges on the calculation of medians across experimental data spans, regardless of the content of those spans, and whether they are readily comparable between recordings, individuals, or conditions. Specifically the process is susceptible to the signal-to-noise ratio (SNR), to the presence of sustained signals, and to the presence of both temporally proximate and distant bursts, and their associated amplitude distributions. To address this, we conducted an examination by simulating data enriched with bursts, elucidating the PT-method’s proficiency in recovering intended bursts and the inadvertent retrieval of unintended ones. Further, and while this also equally applies to the forthcoming issue, we simulated data with conditional differences in power to understand the events recovered in this common experimental scenario. The second issue is that any plausible neurophysiological data will necessarily produce a median and the requisite single-trial maximum correlation, consequently detecting events with a set of characteristics to be classified by their characteristics as either transient or sustained. Hence, the potential of the methodology to erroneously identify transient events within signals of a sustained nature merits consideration. At least three issues of the method may bias our analysis toward a transient interpretation: focusing on the highest-powered data may preferentially capture smaller, sharper events; the use of Morlet wavelet transforms, which weight the center of an oscillatory window more heavily, can amplify brief signal segments; and the reliance on full-width at half-maximum (FWHM) to measure burst duration tends to favor shorter events. Together, these factors could lead to an overrepresentation of transient features at the expense of sustained activity, thereby influencing how we characterize the simulated signals. To confront this issue, we conducted a second simulation, presuming a substrate characterized by both oscillatory and sustained behavior. These analyses focused on the descriptions of event characteristics, with the aim of affirming or negating our assumption of a sustained oscillatory process, based solely on the recovered features.

We establish that the PT-method exhibits robust performance in detecting transient bursts when a relatively high SNR prevails. But its efficacy wanes significantly under conditions of relatively low SNR, leading to the inadvertent detection of numerous undesired events. Furthermore, our examination reveals that when applied to simulations that contain **only** sustained events, the method frequently registers beta events that are clearly within the duration of a burst event, even in cases of a high SNR. That is, if the brain does produce sustained as well as or instead of transient beta events, analyzing them with the PT method will necessarily favor a transient interpretation. Further, we searched across all simulations for datasets that produce a single-trial correlation maximum at six FOM and detailed the extent to which we can differentiate the varied data generation processes based solely on the attributes of the recovered bursts. We conclude that the method has a clearly limited discriminative capacity, in that it consistently yields distributions of burst characteristics that do not easily permit us to draw a distinction between event generation processes. As such determining whether a set of bursts originates from a ground truth process of a burst-filled simulation, a largely noise-filled simulation, or a simulation filled with sustained events is a highly challenging task. Naturally, this lack of discriminative capacity is particularly problematic in the analysis of real neural data, where we have no access to any form of ground truth. Finally, by simulating experimental conditions, we demonstrate how differences in burst counts (or rates) can arise between conditions that differ solely in average power, and have no true rate differences. It is noteworthy that these distinctions appear even when there are no true transient bursts and the oscillatory events are sustained, emphasizing the need for careful interpretation of the method’s results.

## METHODS

### Simulations

We generated two distinct classes of simulations - short-duration wavelet bursts and long-duration sine-wave oscillations - to examine how the PT-method detects transient versus sustained beta activity. Our present simulations were deliberately constrained to scenarios with exclusively transient or exclusively sustained beta, allowing us to probe the PT-method in two extreme regimes. Notably, our low-probability condition does approximate near-complete absence of beta, revealing how the method behaves when bursts are rare.

To create the simulations we embedded either short duration wavelets or long duration sine waves, in continuous 1/f noise, seen in neural recordings, with an exponent of -2 using NeuroDSP function sim_power_law (Cole et al., 2019). The simulations and analyses are all freely available online at https://github.com/langfordzd/burst_simulations. There were 100 trials per simulation, which included 1 second of burst-activity with noise, surrounded by a 1/2 second of pure noise. The sine-waves were asymmetric with rise-decay symmetry uniformly drawn at random between 0.25 and 0.75 using NeuroDSP function sim_cycle. The wavelets were created using the morlet2 function in the python sciPy module (Virtanen et al., 2020) with the whole wavelet being scaled by the amplitude, i.e., the center point of the wavelet was the peak amplitude. In either the wavelet or sine-wave instantiation we randomly flipped the sign of the signal so it would either start with an ascent or descent.The SNR was calculated as the ratio of mean spectral power for the 0.5 seconds in the middle of the trials (the burst-activity period) and 0.5 seconds before the trial (the initial noise period), within the defined beta band.

In these simulations the main parameters of interest were the burst probabilities, the burst durations, and the burst amplitude distributions. The burst probability was manipulated by drawing randomly from a weighted binary distribution at each time-step (250 Hz) to determine if that time-step would start a burst or not – with there being at the least 1 cycle between bursts. The probability of the next time step was specifically weighted to 0.9 in the high, 0.15 in the middle, and 0.03 in the low probability manipulations of the transient event simulations. The probability of the next time step was specifically weighted to 0.9 in the high, 0.1 in the low probability manipulations of the sustained simulations. The burst frequencies (Figure 1B) were always drawn from a truncated normal distribution with a mean of 21, a standard deviation of 1, and being truncated between a low of 15 and a high of 29 Hz. For the transient event simulations we used a lognormal distribution with shape of 0.5, location of 0.04, and scale of 0.12, which we then truncated at 0.4 seconds to model the durations, as shown in Figure 1C. This gave us a distribution similar to empirical research (Shin et al., 2017) that starts around 0.04 seconds, peaks at 0.16 seconds, and tapers off with the highest values somewhere around 0.4 seconds. We simply used a uniform distribution between 0.5 and 0.8 seconds for the durations of the sustained simulations, as seen in Figure 1D. The burst amplitude distributions were modeled again with a lognormal distribution, but we varied the parameters so as to produce significant variation in these amplitude properties and test the effects of the distribution on the outcomes. The impact on amplitude of variations in scale and shape parameters are illustrated in Fig 1A. The location parameter provides a scalar shift along the x-axis of the distribution, in our case therefore a shift in the amplitude. Location was equally spaced in 3 steps between 0.001 and 0.05. The shape parameter is also the standard deviation of the log of the distribution, and affects the general shape of the distribution as seen in Fig 1B. Here it was equally spaced in 9 steps from 0.1 to 0.9. Increasing the scale parameter will spread the distribution along the amplitude axis, and in fact the scale is the median of the distribution. Here the scale parameter was logarithmically spaced in 12 steps between 0.01 and 0.25.

**Figure 1.**
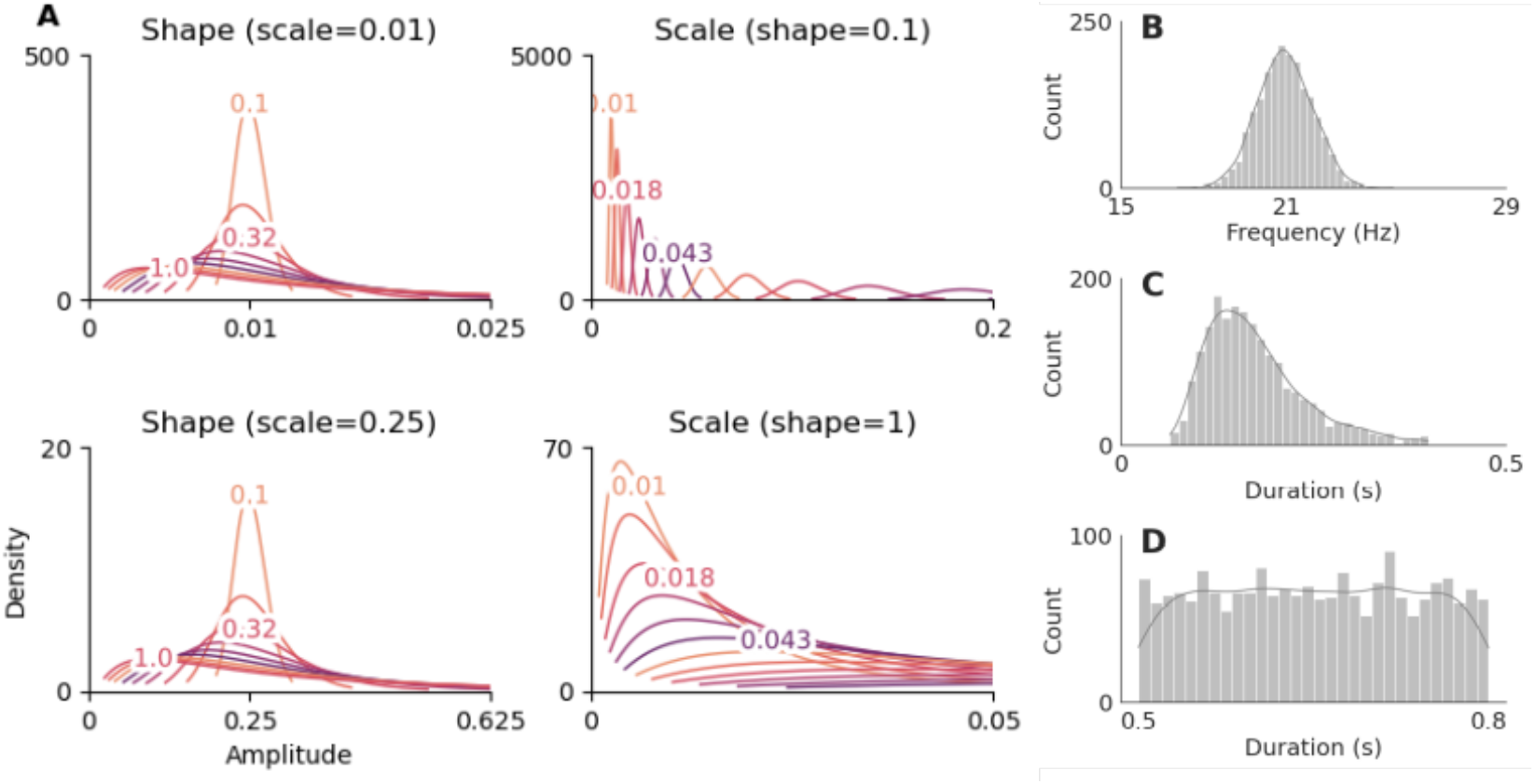
Distributions of simulated burst characteristics. (A) Effects of shape and scale on the amplitude distributions. Representation of the amplitude distributions for each of the shape and scale parameters held constant at the most extreme values of the other (which is indicated in the title of each graph). The location parameter of the distribution was conveniently set to zero, but the location effects on the distribution can be thought as a simple shift along the amplitude axis. **(B) Burst frequency distribution**. Burst frequency was generated from a truncated normal distribution (mean 21 Hz, SD 1 Hz, bounds 15–29 Hz) and remains consistent across amplitude variations. **(C) Transient duration distribution**. Transient burst durations, simulated via a lognormal distribution (shape = 0.5, loc = 0.04, scale = 0.12) and limited to values below 0.4 s, reflect the brief events. **(D) Sustained duration distribution**. Sustained burst durations, drawn from a uniform distribution between 0.5 and 0.8 s, highlight the extended event profile.

#### Rationale for Amplitude, Duration, and Frequency Distributions

We drew on empirical reports (e.g., Shin et al., 2017) indicating that beta bursts in neural recordings often last fewer than 300 ms, prompting our chosen upper duration range for transient bursts. In contrast, our sustained simulations employed oscillations lasting 0.5–0.8 s to represent longer, multi-cycle beta activity that might arise in more tonic neural states. For amplitude, we used a lognormal-like approach (shape, scale, location), inspired by the skewed or ex-Gaussian distributions observed in many oscillatory studies, wherein a sizeable fraction of bursts exhibit relatively modest amplitudes, but there is a long tail toward higher values. Likewise, our beta-band frequency range (15–29 Hz) and distributional choice reflect variability around a singular beta process at 21 Hz that was distributed normally but constrained within the beta band, and would produce a frequency span that has been seen in the literature (Shin et al., 2017).

### Power-threshold detection method

In our instantiation of the PT-method we first translated the simulated signal into a time-frequency representation (TFR) where the signals are convolved with a complex Morlet wavelet at 7 cycles using MNE-Python (Gramfort et al., 2013). The TFR at any time-frequency point was then normalized by the simulation median power value at each frequency; i,e, it was put into factors of medians (FOM). The tested power thresholds ranged from 0.1 to 20 in 30 equally sized steps. For each simulation the best threshold was determined by calculating the maximum correlation between mean single trial power and the area, represented as a proportion of total TFR area per trial, above the cutoff between 15-29 Hz (Shin et al., 2017). Following this, we found local maxima that were above the previously calculated cutoff using skimage function peak_local_max(). These were then considered burst peaks and were used as the dataset to determine the relevant characteristics for each simulation.

We were specifically interested in the power, duration, frequency, and frequency span of all of the recovered bursts. For power we simply took the power value of the detected local maxima, and this was done similarly for frequency. Frequency span was calculated as the full-width half maximum (FWHM) surrounding the local peak in the frequency domain. In a similar vein, duration was calculated as the FWHM surrounding the local peak in the time domain. Following this we determined if each burst was either intended or not, and if more than one detected burst was being assigned to an intended burst we considered it a repeat. A detected burst was considered intended if it had *any* temporal overlap with a simulated burst, but not twice the temporal length. Repeat detections of intended bursts were considered unintended bursts. We then calculated the total number of unintended bursts, the proportion of intended bursts recovered, and importantly for the sustained simulations, the total number of events detected under 0.3 seconds - a duration that we took as a reasonable value to be the transition from transient to sustained events. We then, for each recovered burst, found the maximum absolute value in the time domain signal, and sign flipped it if it was negative. These were then all aligned at the temporal location of the maximum and averaged, which gave us burst related potentials (BURPS; Langford et al., 2023) of each simulation’s collection of detected events.

Beyond the role of the PT-method in our results, the common use of complex Morlet wavelet convolution may bias burst detection by emphasizing the center of the oscillatory window, thereby favoring the extraction of transient bursts even from sustained oscillations. To assess whether this effect is exclusive to Morlet analysis, we conducted a reduced set of simulations (with half the amplitude parameters and fewer trials) using the Stockwell transform—a method with different time–frequency characteristics (Stockwell, et al., 1996). The results (see Supplementary Figure 1) indicate that the Stockwell transform produces similar outcomes, with continuous oscillations being misclassified as bursts.

## RESULTS

We set out to evaluate the PT-method of burst detection using simulated data over a range of plausible parameters of event characteristics. We consider two specific types of simulation. In the first, we’ve assumed that the underlying beta-band process is filled with transient events (0.04-0.3 second durations) and tried to faithfully represent a variety of possibilities close to the burst characteristics that are found in the literature (Morris et al., 2023; Shin et al., 2017). In the second type of simulation we evaluated the results of the PT-threshold method if it were applied to data with a beta-band process that was filled with sustained events (0.5-0.8 second durations), and not transient events. Here, we were interested in what would happen if we incorrectly apply a method to detect transients when they don’t exist. To create the simulations we embedded either short duration wavelets, or long duration sine waves, in continuous fractional noise. In both types of simulations, importantly, we parameterize the amplitudes, the spread of the amplitude distributions, and the probability of oscillatory event occurrence. Taken together, these manipulations inform the signal-to-noise ratio of any given simulation.

### Burst detection assuming that transient events populate the beta-band

The purpose of this simulation was to model the characteristics of bursts found in the literature, assume that they are true, and then see the PT-method’s reaction to different manipulations. Most notably the amplitude distribution and the burst rate are of interest, because in calculating the median cutoffs these should, a priori, have a large effect on the result. We modulated the burst-rate at three levels by weighting the probability of a random draw at each time step (the exact burst count distributions can be seen in column 6 of Figure 3). The burst frequency was assumed to be a truncated normal distribution centralized at 21 Hz with a standard deviation of 1, and truncated between 15 and 29 Hz. The amplitude distribution was a lognormal distribution and we manipulated the location, shape, and scale; the effects of which can be seen in Figure 1A. The SNR then varies, most notably, over the scale parameter of the lognormal distribution, as well as the three steps on the location parameter. The overall duration of the bursts was also set as a lognormal distribution with shape set at 0.5, scale at 0.12, and location set to 0.04. This gave us a distribution similar to empirical research that starts around 0.04 seconds, peaks at 0.16 seconds, and tapers off with the highest values somewhere around 0.4 seconds (Shin et al., 2017). The bursts themselves were modeled as wavelets as this has been the proposed shape in past work (Sherman et al., 2016; Szul et al. 2022; Zich et al. 2023).

In this analysis presented in Fig 2 we see two general trends; the first is that when the amplitude distribution shape is low, across all scales, the method recovers the highest proportion of intended bursts for a given probability distribution. Conversely, as the shape increases the amount of bursts recovered decreases. The second is that as the scale of the distribution increases (increasing the central tendency of the amplitudes) the amount of unintended bursts decreases, for any given shape parameter. Unsurprisingly maybe, this means that as the burst amplitude (SNR) goes up, the chances of an unintended burst goes down, but this relationship is modulated by the spread of the distribution. Going along with this, the maximum correlation threshold both increases as the shape increases, and increases as the scale increases. In Figure 2 the average maximum-correlation threshold at approximately 6 FOM manifests as a diagonal line with a downward-shift in the x-axis, which is the scale parameter of the amplitude distribution. Figure 1 further indicates that the SNR is relatively low when 6 FOM is the correct maximum. In most instances where 6 FOM is applied, the method consistently achieves a high recovery rate, generally exceeding 50%. Higher recovery rates are observed in simulations involving lower probability event detection. A notable challenge lies in the method’s pronounced ability to recover unintended bursts. This phenomenon is evident across all probability levels but becomes more problematic in scenarios with low burst probability. This is attributed to the significantly higher ratio of intended to unintended bursts in such cases.

**Figure 2.**
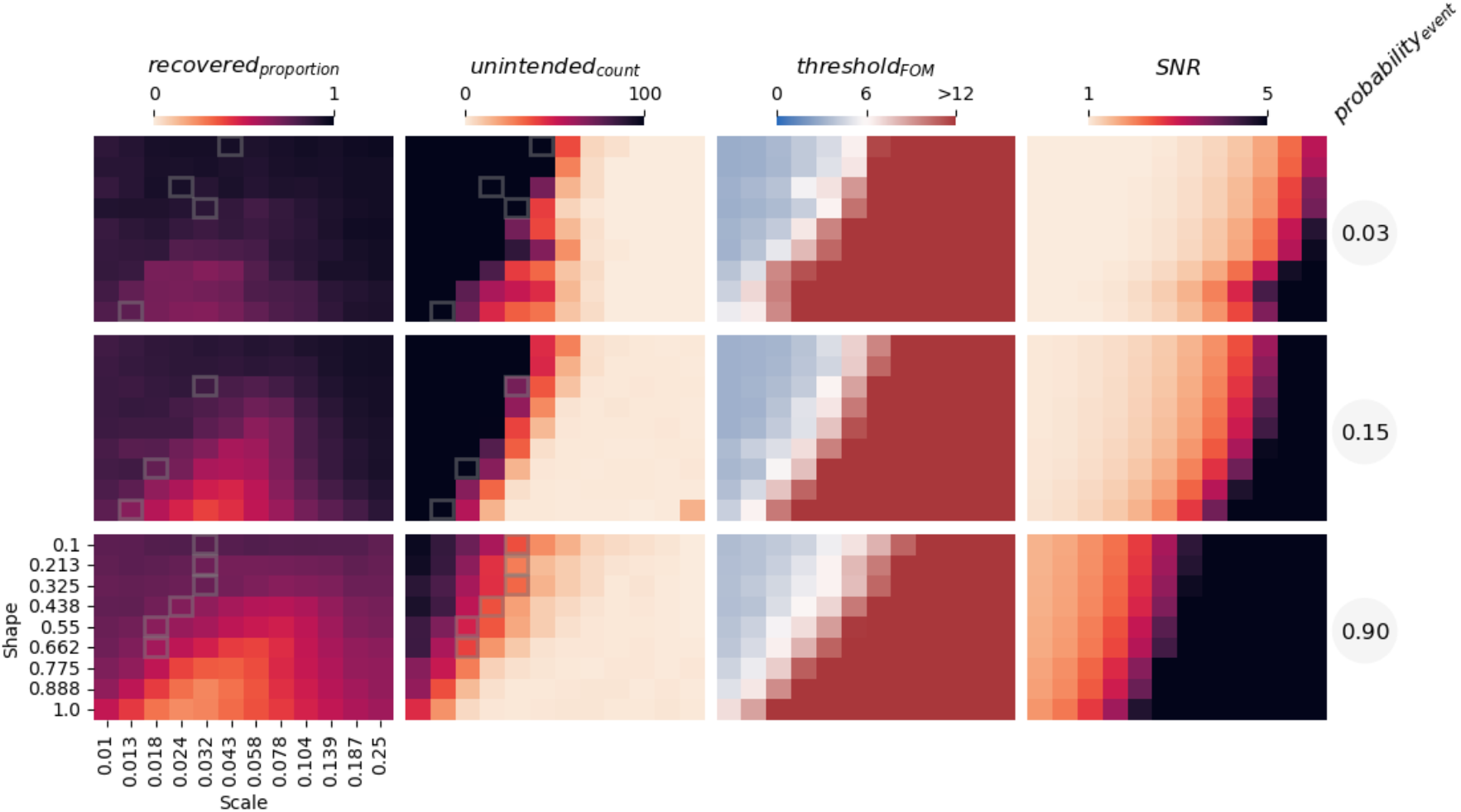
Simulation results when transient events populate the beta band. In each plot, the measure indexed at the top of the column is plotted across variations in shape and scale of the lognormal distribution that has been used to generate burst amplitude. Top row is low event probability, middle is middle event probability, and the bottom is high event probability; all of which are marked on the right of the graphic. Leftmost column is average recovery rate, middle-left is the average unintended count of the 100 trial simulation, middle-right is the average maximum correlation threshold, and the rightmost column is the SNR. Gray boxes indicate points in which the average of the maximum-correlation threshold over the simulations is approximately 6 FOM and indicate where you would most likely find an amplitude distribution that produces this threshold. This demonstrates that even under conditions designed to mimic realistic burst-like activity, threshold-based methods can produce a high rate of unintended detections, especially at lower SNR or broader amplitude distributions.

### Burst detection assuming that sustained events populate the beta-band

In the second type of simulation, we assessed the performance of the PT method when applied to data featuring a beta-band process characterized by sustained events, and not containing the transient events considered above. The key distinction from the transient event simulation lies in the fact that the bursts in this scenario consistently lasted between 0.5 and 0.8 seconds—always sustained. To elaborate, these sustained bursts ranged from a minimum of 7.5 cycles (0.5 seconds at 15 Hz) to approximately 23 cycles (0.8 seconds at 29 Hz), with the duration modeled as a uniform distribution.

For simplicity and to avoid redundancy, we employed only two probability levels, 0.1 or 0.9, recognizing that modeling an even lower probability event would yield results very similar to the lowest probability event in the transient event simulations (i.e., in both there would be very little real beta activity). The frequency distribution mirrored that of the transient simulations. The sustained nature of the signals inherently led to fewer oscillatory events compared to the transient simulation, given temporal constraints.

To address the potential limitations of wavelets for sustained oscillations, we modeled the bursts as sine waves.

As discussed above, real neural signals are reported to contain transient beta bursts, but there is little consideration of whether such transients are the ***only*** instantiation of beta in the signal, or indeed whether more sustained beta processes might be interpreted as transient bursts under certain analysis conditions. As such we wanted to see how the threshold detection method will perform when presented ***only*** with sustained beta events. To test how the method interprets these sustained beta processes, our primary analysis focus shifted from recovering simulated and unsimulated bursts to assessing the characteristic of any recovered bursts that lasted for less than 0.3 seconds—our defined cutoff for a detected event to be considered transient. Correctly identifying sustained events was crucial for evaluating the method’s performance in these circumstances. While recovery of intended sustained events was less relevant to transient burst detection, we didn’t exclude detected sustained events for analysis in the subsequent section.

Figure 3’s leftmost column vividly illustrates that when applying the PT-method to sustained oscillations, it detects <0.3-second transients in the sine wave instantiations, even though these were not included in the simulated data. At the low probability level, the detection pattern on the shape-scale surface closely resembles that of the transient event simulations, reflecting the same minimal oscillatory activity. However, at the high probability level, a distinct profile emerges suggesting that even with a very high signal-to-noise ratio (SNR), the method detects numerous transients.

**Figure 3.**
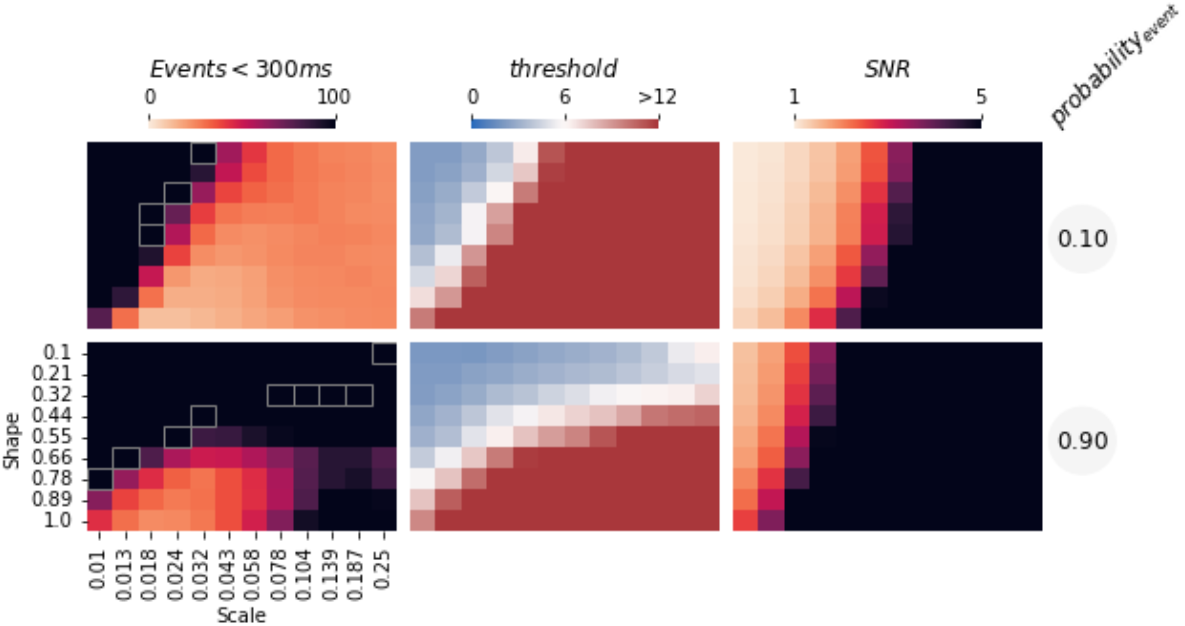
Simulation results when sustained events populate the beta band. Top column is low event probability and the bottom is high event probability. Leftmost column is the count of recovered events that could be considered bursts, the middle is the average maximum correlation threshold, and the rightmost is the SNR. Gray boxes indicate points in which the average threshold is approximately 6 FOM. Thus, these results indicate that purely sustained oscillations can nonetheless be misidentified as short-duration bursts under the PT-method, challenging our ability to distinguish truly transient events from prolonged oscillations.

### Evaluating the characteristics of the recovered bursts

We have therefore demonstrated a capacity of threshold methods to capture unintended bursts in transient simulations and transient events within sustained simulations. As a result, there is an important need for a manner to validate the plausibility of the captured bursts. To do so we now consider the specific characteristics of these bursts captured from the simulated data where we have an established ground truth. Given the prevailing use of a threshold maximum correlation set at 6 FOM in past observations, we focused on simulations where the peak correlation fell within the range of 5.5 to 6.5 FOM.

Figure 4 compiles data from simulations within this peak correlation range, facilitating a qualitative comparison of burst characteristics across various simulated substrates. Our analysis delved into four key characteristics—burst power, frequency span, duration, and burst counts—commonly explored in the literature. Surprisingly, our findings revealed that qualitatively distinguishing between the five types of oscillatory event simulations was not straightforward. Notably, if there is abundant oscillatory signal in the simulation, the median amplitudes of the bursts tend to be higher, even with the same 6 FOM cutoff. This is likely due to the ample signal elevating the median calculation during the FOM procedure and thus possibly contributing to the lower recovery rates in high-probability transient simulations.

**Figure 4.**
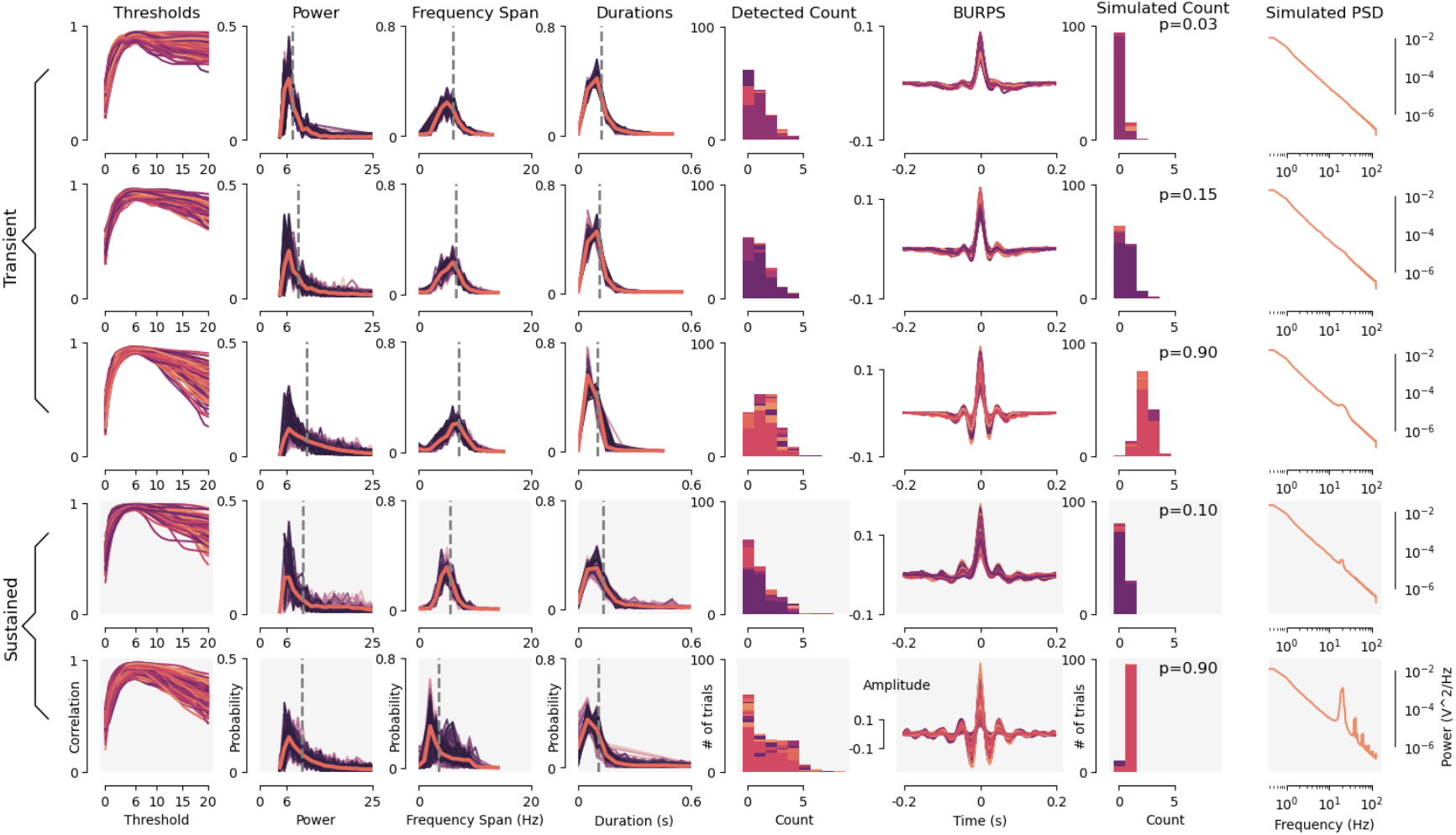
Comparative analysis of burst characteristics across various simulation types, revealing striking similarities among simulation types. Rows 1–3 illustrate transient event simulations set at low (Row 1), mid (Row 2), and high (Row 3) burst probability levels, while Rows 4–5 display sustained event simulations at low (Row 4) and high (Row 5) probability levels. In these simulations, sustained events last between 0.5 and 0.8 seconds, compared to transient bursts, which are modeled as brief events ranging from 0.03 to 0.4 seconds. The first six columns characterize the recovered bursts during the detection process. Specifically, Column 1 shows the correlation at various thresholds with a peak near 6 FOM. Columns 2, 3, and 4 provide the distributions for power, frequency span, and duration of the recovered bursts, with a grey dashed line indicating the median for each distribution. Column 5 reports the trial count, or the number of bursts detected in each dataset, and Column 6 displays the maximum centered burst-related potentials (BURPS). Columns 7 and 8 pertain to the simulated signal. Column 7 presents the intended distribution of trial counts, intended for comparison with the detected count in Column 5, with the event probability marked in the top right corner of each simulated count. Finally, Column 8 depicts the average power spectral density of the simulated signals.

In the frequency domain, we observed minimal differences among simulation types, except in high-probability sustained event simulations, which exhibited a narrower frequency range. This could be explained by fewer bursts overall, albeit with a higher SNR. Interestingly, there was little disparity in the durations of bursts extracted from the different simulations, although sustained simulations appeared to have broader tails in their distributions. Further scrutiny of the threshold correlation profiles and burst count histograms failed to reveal distinct features associating a specific profile with a particular substrate. While some differences were discernible across substrates, a definitive signal of a burst’s origin—from a sustained signal, noise-dominated signal, or one rich in oscillatory signals—remained elusive.

To deepen our analysis, we aligned each detected burst at the largest deflection, incorporating sign-flipping where necessary, and generated averaged burst-related potentials (BURPS) visible in column 7 of Figure 4. BURPS provide a unique perspective on the average rhythmicity of the detected bursts. In both the high-probability burst-filled substrate and the high-probability sustained substrate, the averaged plots displayed a somewhat more rhythmic pattern, characterized by visually more zero-crossings. However, while this rhythmicity provided insight into the generative substrate for each simulation type, again each was subject to be interpreted as having a plausible ‘wavelet-like’ waveform seen in bursts within the broader literature; from very brief ricker-like wavelets (e.g., Sherman et al., 2016) to more temporally extended waveforms (Karvat et al., 2020).

### Differences in burst counts with an average power differential between experimental conditions

Historically oscillatory dynamics have been linked to behavioral or cognitive roles, in part, by considering differences in power between experimental conditions. More recently a similar approach has been applied but using burst count (or similarly burst rate) as a measure instead of power (e.g., Shin et al., 2017). One might consider these two measures to be independent - a count of events, or the power of those events - and a full understanding of the neural dynamics of a system should surely require us to understand their separate role in a system. A potential issue with this approach when working with threshold detection approaches (of whatever form) is that power within transient or sustained events changes the threshold and therefore might change burst detection. In other words, we asked whether burst rate changes could be potentially driven by changes to event power. Our simulation, containing ground truth of how many bursts are inserted into the signal, is a perfect opportunity to test this.

As such, we developed further simulations in which we introduced two simulated behavioral conditions, mirroring situations commonly observed in experimental setups where power differentials exist between conditions. To simulate this, we either shifted the location parameter of the lognormal distributions by a factor of either 1.5, 2, or 3 on half of the trials in the transient or sustained conditions, or simulated half of trials to actually contain more bursts without changing burst amplitude; each of these manipulations create the conditions Low Power (LP) and High Power (HP). We shrank the amount of parameters for the search by only using 3 steps in the shape and scale parameters, and then focused our analysis again at 6 FOM. Specifically, our analysis focused on three distinct scenarios; firstly in conditions abundant with transients (p_event_ = 0.9) and exhibiting a change in event amplitude in half of the trials. Secondly, we examined a scenario featuring abundant (p_event_ = 0.9) sustained events, with half of the trials exhibiting events of higher amplitude. Notably, in neither of these first two simulated scenarios did the event count exhibit systematic changes between conditions, so a perfect detection algorithm should produce identical burst counts in the two conditions, and any detected burst count differences could only be explained by the amplitude differential. Lastly, we simulated a scenario in which the power difference was caused by a transient burst rate difference; in the HP trials the p_event_ = 0.9 and in the LP trials the p_event_ = 0.1. Our attention centered specifically on the transient burst count, given both its significance in numerous studies (e.g., Little, Bonaiuto, Barnes, & Bestmann, 2019; Morris et al., 2023; Shin et al., 2017) and that it indicates the presence of bursts, and thus all other characteristics are dependent on it. Initially, we visualized the single-trial burst count recovered by single-trial power for both scenarios in Fig 5A. The results indicated that as the number of detected bursts increased so did single-trial power, in all scenarios. Consequently, one might anticipate observing condition-wise differences in burst counts in these scenarios, even in the absence of a count differential. Indeed, our findings, depicted in Figure 5B, revealed such differences. When there exist transients there will be more bursts detected in conditions that have relatively higher power caused solely by amplitude differences (Fig 5B top). Remarkably, where there are sustained signals (Fig 5B middle) with a power differential, there will also be a similar finding - a measured difference in burst counts despite there being no such count difference in the simulation. Perhaps unsurprisingly, when the power differential is caused by an actual count difference between conditions, the method detects this count difference (Fig 5B bottom).

**Figure 5.**
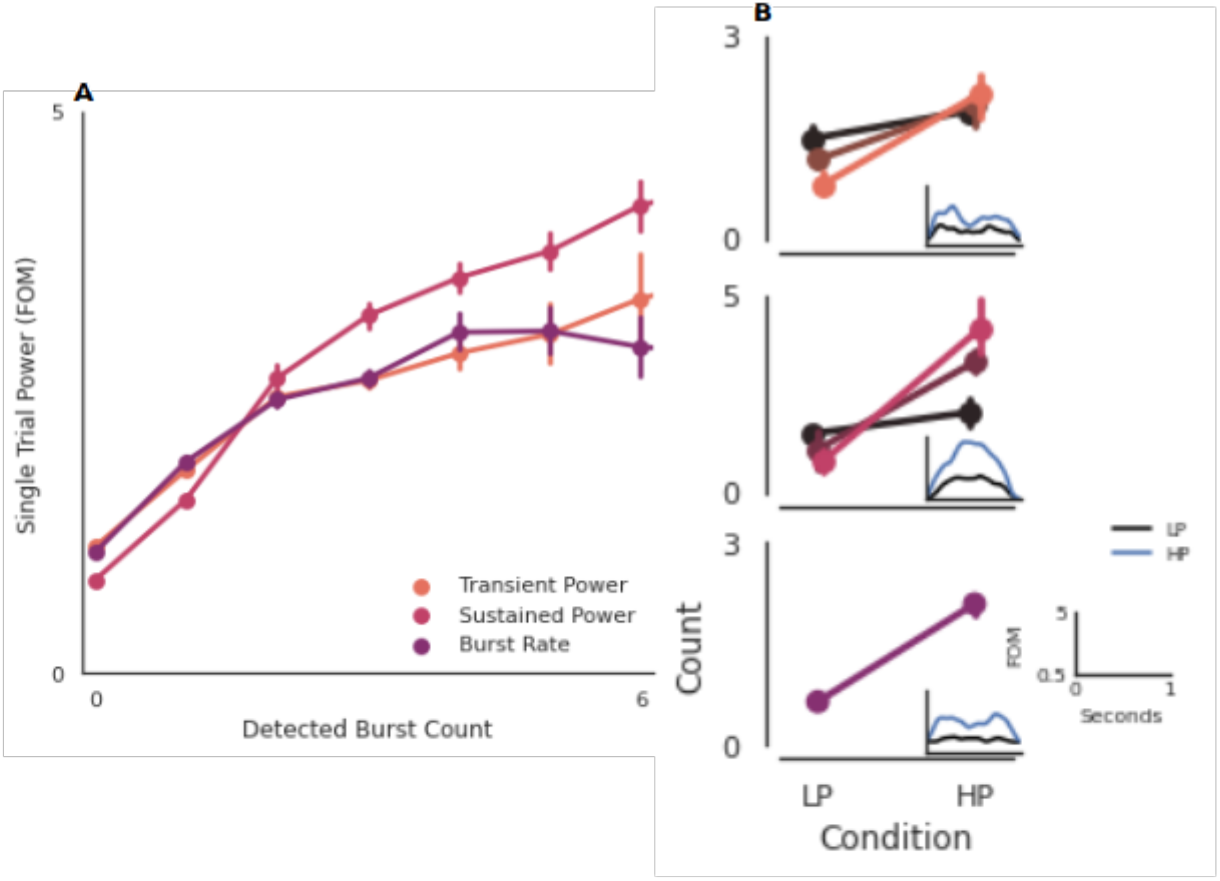
Simulated results highlighting the differences in recovered burst counts in two scenarios where there is no discrepancy in burst counts but a difference in power exists, and one scenario where there is both a burst count discrepancy and a difference in power. (A) Shows the relationship between burst count and power. This indicates a positive correlation between recovered burst count and single trial power across the three types of event simulations driving the power difference between the LP and HP conditions. **(B) Depicts the mean difference in recovered burst count per low and high power conditions for each of the transient (Top), sustained (Middle), and burst rate (Bottom) simulations**. It showcases the averaged burst count detected in the HP and LP experimental conditions. The hues represent the factor by which the location of the amplitude distributions was multiplied to create the HP condition, with the original hue from (A) representing the highest factor and black representing the lowest. The inset demonstrates the average trial power in the HP and LP conditions (using the middle amplitude difference for the transient and sustained). In summary, when power differences arise from actual burst count variations, the method successfully recovers those count differences. Conversely, if the power difference is driven by factors other than burst count, the method may still detect a burst count discrepancy.

## DISCUSSION

Here we have assessed the effectiveness of the widely used thresholding method in detecting beta bursts using simulated neural signals. We manipulated the SNR by varying burst-rate probability and burst amplitudes in simulated data. We conducted two distinct types of simulation, each based on different assumptions. In the first type, where a “bursty” substrate was assumed, we examined the PT-method’s ability to recover intended bursts while also evaluating its capacity to identify unintended bursts. The method’s performance hinged on the shape and scale of amplitude distributions, along with the prevalence of transient events. The method recovers intended events at a reasonable rate, yet is prone to detecting an over-abundance of unintended events. In the second type of simulation, assuming a substrate without bursts but featuring sustained oscillations, our results were more concerning. Even in cases of clearly prolonged events and a high SNR, the method demonstrated a tendency to recover numerous short-duration events that could be misinterpreted as bursts. Subsequent analysis of each simulation type emphasized that while the PT-method may perform well under specific conditions, differentiating between bursty signals, predominantly colored noise, and sustained signals based on the recovered characteristics presents a considerable challenge. In our simulations of experimental conditions, we reveal how variations in burst counts can emerge solely due to differences in average power. Notably, these distinctions persist even during sustained oscillatory events emphasizing the potential deceptive nature of behavioral correlations. Together, this challenges the reliability of research utilizing the PT-method and urges a reevaluation of its application.

Given the challenges and limitations associated with the PT-method in accurately identifying and characterizing beta bursts, it is crucial to complement the analysis with analysis of behavioral correlates. Though, as we highlight here, this is not without its own complications. In instances where the method detects bursts that coincide with behavioral events, investigating the precise temporal dynamics and examining the relationship between burst occurrence, burst properties, and behavior could provide meaningful insights into cognitive processes. Yet, it is important to remain cautious and consider alternative explanations. As we indicate here, when there exists a difference in power across conditions, and a burst rate difference is also detected, it is completely feasible that such a rate difference is driven by other factors that can influence power, and not by burst rate. And this distinction is critical for any mechanistic interpretation of the role of oscillations within a network. If a power difference exists between conditions, it is important to be able to interpret with confidence the source of that difference - if the difference comes from changing burst rates, that gives us specifically and potentially mechanistic information about the network dynamics, an interpretation that may well not hold if the power differences are driven by separate factors. Furthermore, it is possible that the broader beta power dynamics could exhibit temporal patterns that may influence burst detection and apparent associations with behavior, e.g., it is known the beta power can modulate consistently over entire recording sessions (Stoll et al., 2016). Such temporal dynamics could introduce complexities in the interpretation of burst-behavior relationships, highlighting the importance of carefully considering the broader context and potential confounding factors.

Crucially, even when the PT-method identifies bursts strongly correlated with behavior and there exists no power differential in the conditions, asserting a high degree of confidence in characterizing the phenomenon as transient, rather than sustained, is unwarranted. Simply pinpointing these bursts doesn’t guarantee their transient nature. Realistic neural signals inherently exhibit a median and corresponding single-trial maximum correlation, resulting in events that might superficially appear transient when thresholded – a tendency observed in many cases. This issue stems from imperfections in the PT method. For sustained signals, setting a threshold too high may capture only the most powerful segment, potentially overlooking the signal’s sustained nature. Essentially, the most powerful segments, often of shorter duration, excel at explaining variance in single-trial power compared to lower power and longer segments in a neural signal; and as such the threshold is set to high. In this context, we posit that the PT-method coerces numerous realistic scenarios into being incorrectly interpreted as being populated by transient events - a point overlooked in the broader literature. Therefore, before placing confidence in its outcomes, it becomes crucial to establish the veracity of the assumption that the data indeed comprises transient events rather than sustained ones.

Moreover, while we have focused primarily on how the method can inflate or mask differences in burst rate, it may likewise affect the *timing* of detected bursts. Setting a threshold that is too high or low could artificially shift the onset or offset of detected bursts in the time–frequency map similar to burst-rate, potentially creating misalignment between true neural events and behavioral markers. This concern is particularly salient given findings that *when* a burst occurs (i.e., burst timing) can be more predictive of behavior than average burst power (Little et al., 2019). Consequently, future work should investigate how threshold-based methods might alter burst timing under various signal-to-noise conditions, and how these distortions could influence correlations with behavioral or cognitive outcomes.

Acknowledging these limitations underscores that a critical re-evaluation of existing literature on beta bursts is essential, involving a meticulous examination to determine the transient nature of the phenomenon. This reassessment serves as a foundational step toward refining our knowledge of the beta band and its temporal dynamics. Further, it is likely necessary for a comprehensive investigation to be based on multiple methods. The utilization of diverse and complementary techniques, such as elaborating on past research using hidden Markov models (e.g., Heideman et al., 2020); including recent time-delayed embedded models, e.g., (Rier et al., 2024)), using lagged-coherence to detect bursts or verify rhythmicity before burst detection (Fransen, et al., 2015; Rayson et al., 2022), adapting the cycle-by-cycle method (Cole et al., 2019; e.g., Langford et al., 2023), or methods incorporating 1/f corrections (e.g., Rodriguez-Larios et al., 2023; Szul et al., 2023; Brady et al., 2022) offer intriguing alternatives for isolating beta events from background noise, and might facilitate the cross-validation of burst detection results and offer a more robust insight into the underlying neural dynamics.

## Supplementary Materials

**Supplementary Figure S1.**
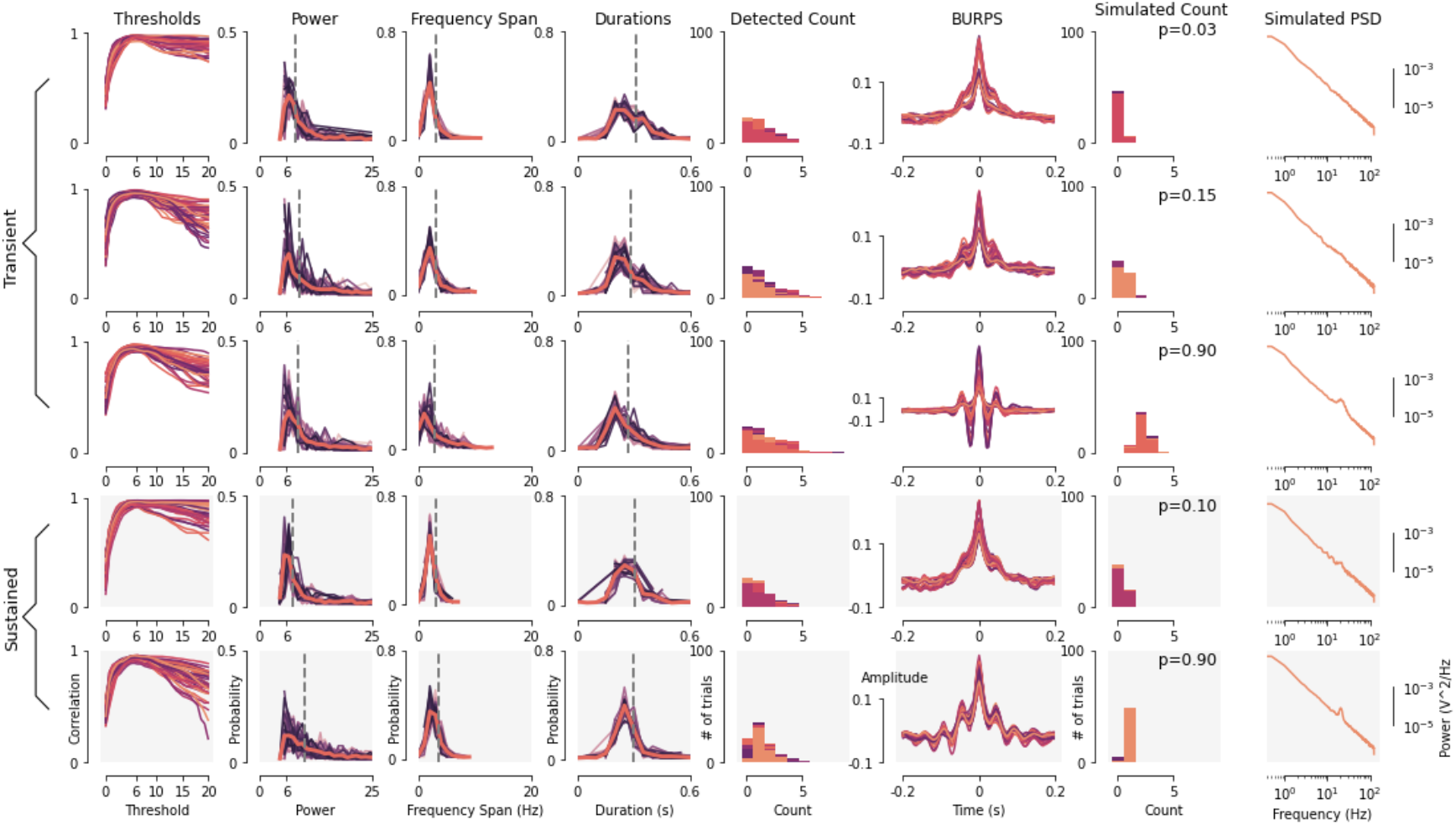
Analysis of burst characteristics using the Stockwell transform and a reduced simulation. Rows 1–3 illustrate transient event simulations at low, mid, and high burst probability levels, whereas Rows 4–5 display sustained event simulations at low and high probabilities, respectively. In these simulations, transient bursts are modeled as brief events (0.03–0.4 s) and sustained events range from 0.5 to 0.8 s. The first six columns characterize the detected bursts: Column 1 shows the correlation across thresholds (peaking near 6 FOM), Columns 2–4 present the distributions for burst power, frequency span, and duration (with dashed lines indicating the medians), Column 5 reports the trial count, and Column 6 displays the maximum burst-related potentials (BURPS). Columns 7 and 8 pertain to the simulated signal—with Column 7 depicting the intended trial count distribution (with event probability indicated in the top right corner) and Column 8 showing the average power spectral density. Despite employing the Stockwell transform in this reduced simulation, the results indicate that sustained oscillations can still appear as bursts, suggesting that the artifactual detection of bursts is not solely attributable to the use of Morlet wavelet convolution.

